# Use of CRISPR interference for efficient and rapid gene inactivation in *Fusobacterium nucleatum*

**DOI:** 10.1101/2023.09.19.558491

**Authors:** Peng Zhou, Bibek G C, Flynn Stolte, Chenggang Wu

**Affiliations:** Department of Microbiology & Molecular Genetics, the University of Texas Health Science Center, Houston, TX, USA

**Keywords:** *Fusobacterium nucleatum*, CRISPRi, Gene inactivation, Riboswitch, essential gene

## Abstract

Gene inactivation via creating in-frame deletion mutations in *Fusobacterium nucleatum* is time-consuming, and most fusobacterial strains are genetically intractable. Addressing these problems, we introduced a riboswitch-based inducible CRISPRi system. This system employs the nuclease-inactive *Streptococcus pyogenes* Cas9 protein (dCas9), specifically guided to the gene of interest by a constantly expressed single guide RNA (sgRNA). Mechanistically, this dCas9-sgRNA complex serves as an insurmountable roadblock for RNA polymerase, thus repressing the target gene transcription. Leveraging this system, we first examined two non-essential genes, *ftsX,* and *radD*, pivotal for fusobacterial cytokinesis and coaggregation. Upon adding the inducer, theophylline, *ftsX* suppression caused filamentous cell formation akin to chromosomal *ftsX* deletion, while targeting *radD* significantly reduced RadD protein levels, abolishing coaggregation. The system was then extended to probe essential genes *bamA* and *ftsZ*, vital for outer membrane biogenesis and cell division. Impressively, *bamA* suppression disrupted membrane integrity and bacterial separation, stalling growth, while *ftsZ-*targeting yielded elongated cells in broth with compromised agar growth. Further studies on *F. nucleatum* clinical strain CTI-2 and *Fusobacterium periodonticum* revealed reduced indole synthesis when targeting *tnaA*. Moreover, silencing *clpB* in *F. periodonticum* decreased ClpB, increasing thermal sensitivity. In summary, our CRISPRi system streamlines gene inactivation across various fusobacterial strains.

**IMPORTANCE:** How can we effectively investigate the gene functions in *Fusobacterium nucleatum*, given the dual challenges of gene inactivation and the inherent genetic resistance of many strains? Traditional methods have been cumbersome and often inadequate. Addressing this, our work introduces a novel inducible CRISPRi system in which dCas9 expression is controlled at the translation level by a theophylline-responsive riboswitch unit, and sgRNA expression is driven by the robust, constitutive *rpsJ* promoter. This approach simplifies gene inactivation in the model organism (ATCC 23726) and extends its application to previously considered resistant strains like CTI-2 and *Fusobacterium periodontium*. With CRISPRi’s potential, it is a pivotal tool for in-depth genetic studies into fusobacterial pathogenesis, potentially unlocking targeted therapeutic strategies.

## INTRODUCTION

*Fusobacterium nucleatum* is an anaerobic, Gram-negative bacterium, often found in modest quantities within health subgingival dental plaque (1, 2). However, its numbers surge in periodontal pockets (3–5). Beyond its association with periodontitis, significant evidence suggests that when *F. nucleatum* spreads beyond the oral cavity, it is implicated in numerous systemic ailments, including colorectal cancer, adverse pregnancy outcomes, inflammatory bowel disease, and rheumatoid arthritis (6, 7).

This bacterium displays remarkable genotypic and phenotypic diversity. Currently, it’s divided into four subspecies: *nucleatum*, *vincentii*, *animalis*, and *polymorphum* (8, 9). Genome analyses have revealed unique gene clusters in each subspecies, emphasizing their considerable genetic variations (10–12). Some researchers believe these differences warrant a reclassification of these subspecies into a higher taxonomic rank (13). These genetic discrepancies may influence their varied behaviors, such as adaptability to multi-species biofilms (14), the formation of single-species biofilms (15), and interactions with neutrophil functions (16). In addition, these variations might be the foundation of their specific roles within the human body. For instance, *nucleatum* is associated mainly with periodontal disease (17), *animalis* is linked with colon-related disorders (18, 19), and *polymorphum* is tied to pregnancy complications (20). Notably, *vincentii* often coexist with *polymorphum* in healthy mouths, typically seen as benign (17). Yet, recent insights suggest its potential role in prostatic inflammation, hinting at connections to prostate disorders (21).

Beyond this, even within the subspecies, genetic variations abound. For example, consider three strains of subspecies *nucleatum*: ATCC 25586, ATCC 23726, and CTI-2. ATCC 25586, isolated from a cervicofacial lesion, became a touchstone in early fusobacterial research, shedding light on *F. nucleatum*’s nutrition, physiology, genetics, and biochemistry. In liquid cultures, its cells tend to cluster and settle, often growing beyond 4 microns (22), and challenges arise when attempting to transform this strain using shuttle plasmids (23, 24). These set it apart from the *F. nucleatum* strain ATCC 23726, sourced from the human urogenital tract. Typically, its cells measure between 1-2.2 microns and do not self-aggregate under standard conditions (25), and its standout traits are a robust transformation capability and genetic tractability (26). This has positioned ATCC 23726 as a favored model organism for studying *F. nucleatum* virulence factors. The transformation disparities between ATCC 25586 and ATCC 23726 can be attributed to variations in their restriction-modification systems (23) – such variations that hinder most fusobacterial strains from tapping into their full genetic potential. Then there’s CTI-2, another strain within the subspecies *nucleatum*, originating from human colorectal cancer samples (27, 28). Among its counterparts, CTI-2 stands out due to its genome housing an intact operon encoding a type IV secretion system (T4SS) – a feature missing in ATCC 25586 and 23726 sequenced genomes. Yet, it’s prevalent in the subspecies *animalis* and *polymorphum* strains (https://img.jgi.doe.gov/). T4SS, pivotal in bacterial conjugation, DNA exchange, and pathogenesis (29), underscores the inconsistency in the distribution of some virulence genes across subspecies of *F. nucleatum*.

Past studies on *F*. *nucleatum*-associated disease mechanisms primarily revolve around ATCC 23726 (25, 30–34) and strain 12230 (35, 36), with the latter belonging to the subspecies *polymorphum*. However, given the diversity of *F. nucleatum*, focusing solely on these two domesticated strains could lead us to miss other significant behaviors and features of this bacterium. This limited focus hampers our holistic understanding of its physiological and pathological aspects. To truly delve into *F. nucleatum*’s intricate nature and disease-causing capabilities, broadening our research to include a range of strains is essential, setting the stage for better clinical practices and treatment methods.

Yet, this endeavor isn’t without challenges. Most fusobacterial strains resist current gene inactivation methods, which need to introduce a suicide plasmid into cells (26, 35, 37, 38). This plasmid, designed to target and disrupt a specific gene, requires either a segment homologous to a portion of the target gene or two segments that border the target site on the host chromosome. Once inside, the plasmid might be recombinant with the host chromosome at the homologous sequence(s), potentially leading to the inactivation or replacement of the target genes. *F. nucleatum* presents a hurdle: its recombination efficiency is at a paltry ∼0.05% (39). Consequently, the demand for a high transformation rate becomes imperative to achieve any meaningful recombination. Unfortunately, most fusobacterial strains exhibit minimal transformation propensity, primarily due to intricate and diverse restriction-modification systems (23). For instance, introducing 1 µg shuttle plasmid pCWU6 into ATCC 23726 can yield over 10,000 transformants, but for ATCC 25586, the same quantity of plasmid only results in less than 10 transformants (24, 38). The Slade lab has found a workaround by employing host DNA methyltransferase to modify the plasmid before electroporation, significantly boosting the transformation rate in ATCC 25586 and enabling gene deletions (23). Regrettably, this modified plasmid system only works for this strain and not for other hard-to-genetically modified strains, restricting its applicability. Another challenge lies in our two-step approach for creating an in-frame deletion in ATCC 23726; it is a time-intensive method (25, 32, 39). This is especially true when investigating essential genes (40), which are particularly interesting to researchers. These genes often underpin critical cellular processes and present promising targets for antimicrobial drug development.

In light of these challenges, we turned our attention to the CRISPR interference (CRISPRi) technique, which bypasses the need for homologous recombination (41). This recently advanced method has proven its merit in studying gene functions across many bacterial species (42, 43). At its core, the CRISPRi system operates with two main components: the dCas9, a nuclease-inactive variant of *Streptococcus pyogenes* Cas9 protein, and the single guide RNA (sgRNA). This sgRNA is crafted with a 20-nucleotide (nt) specific complementary segment with the target gene (base-pairing region), a 42-nt section that binds to Cas9 (dCas9 handle), and a 40-nt transcription terminator originating from *S. pyogenes* (41). When the sgRNA pairs with a gene’s non-template strand, the resulting dCas9-sgRNA-DNA complex blocks RNA polymerase (RNAP), inhibiting the transcription of the target gene (Fig.1A).

**Figure 1:**
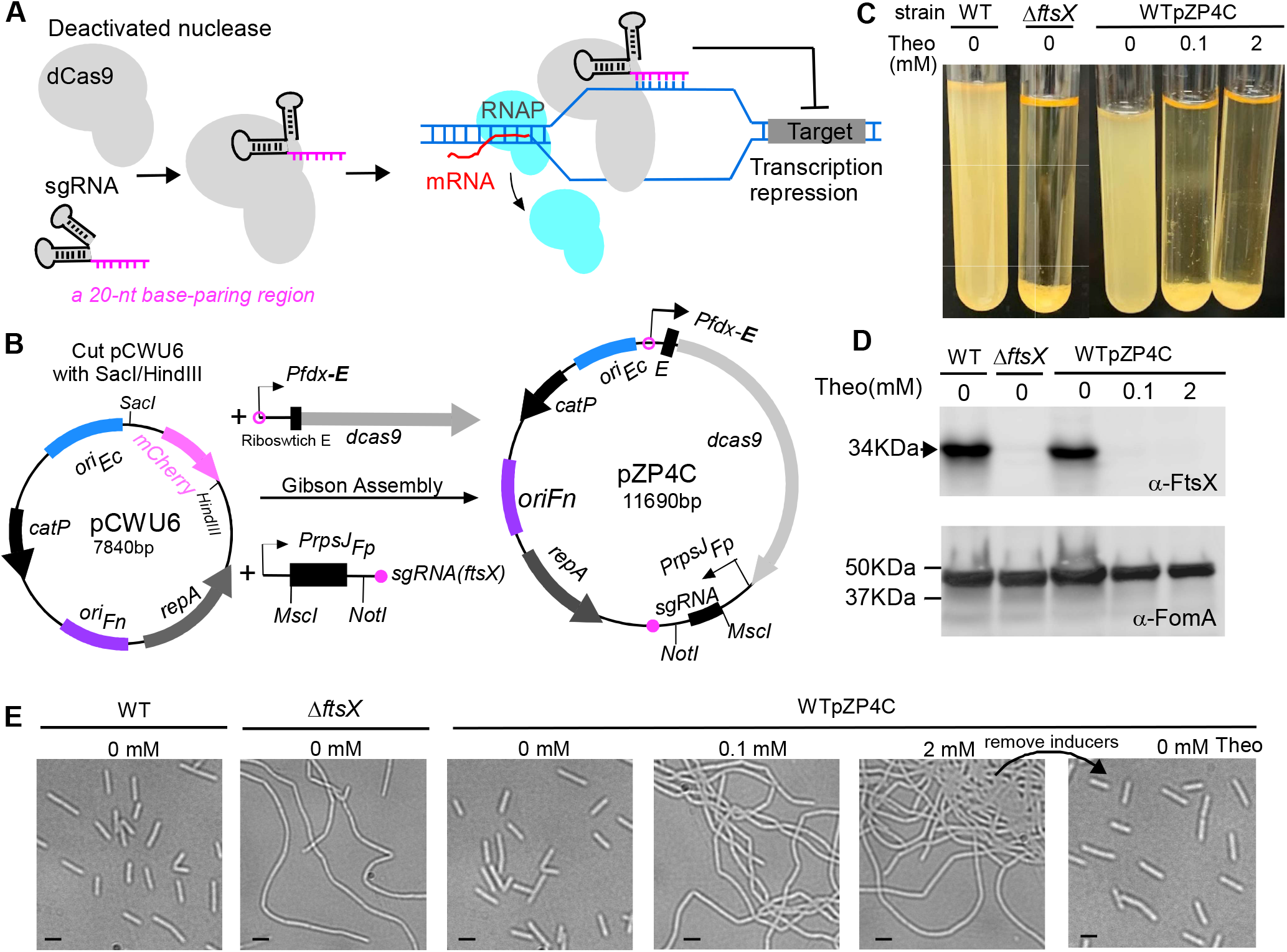
Construction and characterization of CRISPRi system in *Fusobacterium nucleatum*. **(A)** Mechanism of CRISPRi. The nuclease-deficient (dCas9) binds single guide RNA (*sgRNA*) to target the gene of interest via a 20-nt base-paring region. Instead of cleaving DNA, dCas9 obstructs RNA polymerase (RNAP) movement and inhibits transcription elongation, suppressing the target gene’s expression. **(B)** pZP4C construction Blueprint for CRISPRi. pZP4C is derived from the *E*. *coli-F. nucleatum* shuttle vector pCWU6 and is aimed at *ftsX*. pCWU6 underwent SacI/HindIII digestion and was conjoined with two PCR fragments (P*fdx*-*E-dcas9* and P*rpsJ_FP_*-*sgRNA*) through Gibson assembly, resulting in pZP4C. Within pZP4C, the *dCas9*’s expression is governed by P*fdx* and a theophylline-responsive riboswitch E (indicated “E”), both at the transcriptional and translation stages. The *sgRNA* is driven by a constitutive active P*rpsJ* promoter from *Fusobacterium periodonticum* and can be replaced using the MscI and NotI restriction sites. Feature depicted: The P*fdx-E* is sourced from pREcas1, a theophylline-regulated gene expression vector for *Clostridioides*. The *dCas9* and *sgRNA* element originates from pIA33, a *Clostridioides difficile* CRISPRi plasmid; *oriFn* and *repA*, the replication region; *oriEc*, replication region of the *E. coli* plasmid pBR322; *catP*, the chloramphenicol acetyltransferase gene, conferring resistance to thiamphenicol in *F. nucleatum* or chloramphenicol in *E. coli.* **(C)** *ftsX* gene suppression via CRISPRi. The wild-type *F. nucleatum* strain with pZP4C was cultivated in a TSPC medium under varying theophylline concentrations (0, 0.1, and 2 mM) for 22 hours anaerobically. Induction of *dcas9* expression by theophylline in the wild-type cells containing pZP4C results in cells setting at the tube’s base after overnight growth, leaving a transparent supernatant characteristic similar to the *ftsX-*deletion phenotype. **(D)** Western blot analysis validates *ftsX* silencing. Cell samples equivalent to those in panel (C) alongside an additional control sample from the wild-type without inducer, were subjected to SDS-PAGE and then immunoblotted. Antibodies specific for FtsX (α-FtsX) and FomA (α-FomA) were used, with the latter as a loading control. The positions of molecular mass markers (in kilodaltons) are indicated to the left of the blot. **(E)** Silencing FtsX alters fusobacterial cell morphology. A Phase-contrast microscopy recorded cell shapes in panel (B). Bars, 2 µm.

For this study, we constructed a pCWU6-based (25) plasmid equipped with a CRISPRi system. Within this system, the dCas9 expression is governed by a riboswitch-controlled inducible promoter, while the *rpsJ* promoter manages sgRNA’s expression. We showcased its effectiveness by deactivating/knocking down two non-essential genes (*ftsX* and *radD*) and two essential genes (*bamA* and *ftsZ*) in model organism ATCC 23726. Furthermore, we applied this method to examine gene functionality in the highly resistant clinical strain CTI-2 and *F. periodonticum.* CRISPRi presents a faster avenue for gene inactivation than the homologous recombination-based method in *F. nucleatum*, offering significant advantages when investigating essential genes and resistant strains.

## RESULTS

### Construction of a riboswitch-controlled CRISPRi system for repressing gene expression in *F. nucleatum*

To manipulate gene expression in *F. nucleatum* using the CRISPRi technique, we employed the fusobacterial replicable vector pCWU6 (25) as a backbone, creating the pZP4C plasmid via a Gibson assembly procedure (Fig.1B). This pZP4C incorporates the CRISPRI system’s two key components: *dcas9* and *sgRNA* (Fig. 1A & 1B). The chosen *dcas9*, sourced from *S. pyogenes* and codon-optimized for *C. difficile*, is expected to function optimally in *F. nucleatum*, given their similar low GC content. In pZP4C, the synthetically regulatory element *Pfdx-E* (44) controls the *dcas9* expression, composed of the *fdx* promoter and a theophylline-responsible riboswitch E unit (“E” in Fig. 1B). Originating from the *Clostridioides sporogenes* ferredoxin gene Clspo_c0087, the *fdx* promoter partners with the riboswitch – an aptamer and a synthetic ribosome binding site fusion – to tune target gene expression translationally finely (45). Meanwhile, the strong, constitutive *rpsJ* promoter from *F*. *periodonticum* ATCC 33693 (39) drives sgRNA expression.

Our designed sgRNA in pZP4C features a 20-nt base-pairing region targeting *ftsX*, a gene pivotal for fusobacterial cell division. Deletion or depletion of *ftsX* induces morphological changes in broth-grown cells (39, 40): they become elongated, intertwine, and form aggregates that settle at the culture tube’s bottom, leaving the supernatant clear (Fig. 1C). In contrast, wild-type cells lead to a uniformly turbid medium (Fig. 1C). Such marked phenotypic shifts from *ftsX* alternations offer a direct means to assess the CRISPRi system’s impact on *F. nucleatum*. To evaluate our CRISPRi system’s efficiency, we introduced the pZP4C plasmid into the model strain for fusobacterial genetic studies, ATCC 23726. This produced the WTpZP4C strain. When cultivated in TSPC media with varying theophylline concentration, this strain reveals a dCas9 expression profile that we surmise aligns with the inducer concentration. With the guidance of a constitutively expressed sgRNA, the expressed dCas9 binds to the *ftsX* target site, establishing a transcriptional block. Thus, high inducer dosages for an operational CRISPRi system should effectively diminish *ftsX* expression, mirroring the sedimentation observed in *ftsX*-deleted mutants. True to our hypothesis, WTpZP4C cells with 2 mM theophylline mirrored the sedimentation patterns of *ftsX* deletion mutants (Fig. 1C). Remarkably, a minimal 0.1 mM theophylline prompted similar behavior (Fig. 1C). Conversely, without the inducer, the WTpZP4C cells exhibited growth dispersion, like wild-type strains, reflected in their turbidity (Fig. 1C).

Considering potential off-target effects inherent to some bacterial CRISPRi systems, it was crucial to validate that the observed aggregation phenotype stemmed from *ftsX* suppression. We subjected WTpZP4C cells, grown under different theophylline concentrations, to Western blotting, probing with an FtsX-specific antibody. The analysis unambiguously indicated the absence of FtsX protein in cells exposed to both 0.1-and 2-mM theophylline (Fig. 1D). These conditions drove the cells into a filamentous form (Fig. 1E). Without the inducer, pZP4C cells displayed FtsX levels consistent with wild-type strains (Fig. 1D & E), underscoring our CRISPRi system’s precision and efficacy. We removed the inducer from the WTpZP4C culture to verify the reversibility of CRISPR-mediated repression. The elongated cells returned to an indistinguishable morphology from wild-type cells (Fig. 1E, last panel). Overall, our finding robustly demonstrates the proficiency of the riboswitch-controlled CRISPRi system in gene repression within *F. nucleatum*.

### mCherry reporter-assisted sgRNA cloning for efficient CRISPRi plasmid construction

To bolster the efficiency of this system, we developed a streamlined method for generating CRISPRi plasmids. In the pZP4C, two restriction sites MscI and NotI flank sgRNA(*ftsX*), designated for cloning new sgRNAs (Fig. 2A). Each sgRNA consists of a 20-nt base-paring region, a 42-nt dCas9 handle, and a 40-nt transcription terminator (Fig. 2A). For every new CRISPRi plasmid, only the 20-nt base-paring segment in the sgRNA needs to be changed, which is achievable by modification at the center of primer P1 (Fig. 2A and Table 2). Notedly, P1 has three sections: a 21-nt *rpsJ* promoter sequence for future Gibson assembly, an adjustable 20-nt base-paring region at the center, and a 19-nt segment that aligns with the dCas9 handle of sgRNA. When P1 and P2 (Fig. 2A) are used for PCR with pZP4C as the template, it produces a 196-bp DNA fragment, including a new sgRNA targeting the new gene. This fragment can be integrated into the MscI and NotI-digested pZP4C using Gibson assembly to generate a new CRISPRi plasmid, pZP4C(*new*). However, the double enzyme digestion of pZP4C often leads to inconsistent results, producing a mixture of fully digested, singly digested, and undigested plasmids. Isolation of fully digested plasmids on a DNA agarose gel becomes challenging due to size similarities. Single digestion can cause self-ligation, together with undigested plasmids, producing a substantial background when transformed into cloning *E. coli*. This demands colony PCR to identify positive clones, but given the minor 20 nucleotide difference between the new and parent plasmids, differentiation becomes a challenge, often necessitating DNA sequencing. To streamline cloning, we introduced pZP4C-*mCherry* (Fig. 2A), incorporating a *mCherry* reporter cassette (P*rpsJ_Cd_*-*mCherry*) between MscI and NotI of pZP4C. This cassette yields a vivid *mCherry* expression, providing a visual marker to identify undesirable ligation products - colonies containing pZP4C-*mCherry* glow red, simplifying the identification of desired clones.

**Figure 2:**
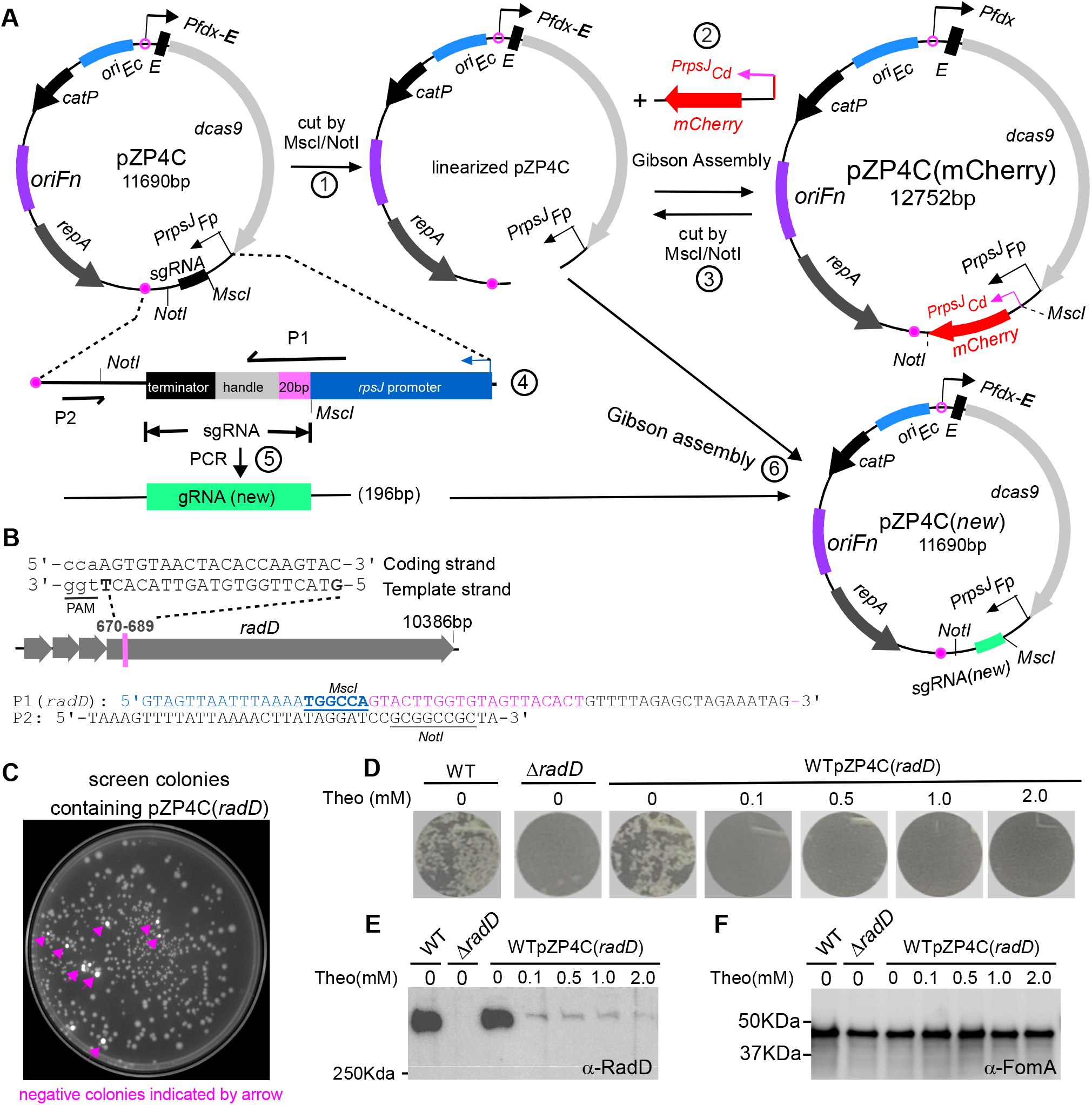
Use of mCherry as a reporter for streamlined screening of positive CRISPRi clones and exemplifying CRISPRi targeting with *radD*. **(A)** Diagram illustrating the construction of CRISPRi plasmid with mCherry as a screening reporter is outlined.1. Linearization of pZP4C: The pZP4C plasmid is cleaved using MscI and NotI enzymes. 2. Replacement of sgRNA segment with the *mCherry* gene. The *mCherry* is driven by the P*rpsJ_Cd_* promoter from pCWU6. The resulting plasmid is designated as pZP4C(*mCherry*), which is then utilized for introducing the new gene target. 3. Release of P*rpsJ_Cd_* -*mCherry*: pZP4C(*mCherry*) is again cut using MscI/NotI to release P*rpsJ_Cd_* -mCherry for subsequent cloning. 4. Design Primers P1 and P2. Primer P1 is meticulously designed in a 5’ to 3’ direction. It begins with a 21-nt sequence matching the P*rpsJ_FP_* promoter region in pZP4C, followed by a 20-nt sequence tailored for the target gene, and concludes with a 19-nt segment targeting the *sgRNA* handle region. The *sgRNA* includes a 20-nt base-pairing region, a 42-nt dCas9-binding RNA structure (dCas9 handle), and a 40-nt transcription termination sequence from *S. pyogenes*. 5. Amplification of new target *sgRNA* fragment: using pZP4C plasmid DNA as a template, primers P1 and P2 amplify a 196-bp new target *sgRNA* fragment. 5. Cloning of new *sgRNA*: The new *sgRNA* fragment is cloned into digested pZP4C(mCherry) using Gibson assembly, yielding a new CRISPRi plasmid targeting a new gene. **(B)** Designing P1 primer with *radD* gene example. For instance, the *radD* gene illustrates the P1 primer design. A 20-nt base pair region from the 670^th^ to the 789^th^ base of *radD* is selected. This sequence, aligned in a 5’ to 3’ orientation on the template strand, is incorporated into the P1 primer (designated P1(*radD*)). Notably, the PAM (TGG) sequence adjacent to the target region is indicated through underlining in lowercase. The MscI site in P1 and the NotI site in P2 are shown. **(C)** Positive clone selection uses a ChemiDoc^TM^ MP Imaging System (Bio-Rad). Fluorescent cells, considered negative clones, appear pseudocolored in white under the Cy3 setting, whereas non-fluorescent cells (positive clones) are gray. **(D)** CRISPRi-mediated repression of the *radD* gene abolishes coaggregation with *A. oris*. The wild-type *F. nucleatum* strain containing pZP4C(*radD*) was cultured in a TSPC medium under different theophylline concentrations (0, 0.1,0.5,1, and 2 mM) for 22 hours in anaerobic conditions. Subsequently, cells were collected, washed, resuspended in a coaggregation buffer, and assessed for coaggregation with *A. oris*. A representative result is presented after three experimental repeats. **(E and F)** cells subjected to the experimental conditions detailed in panel (D) undergo SDS-PAGE followed by immunoblotting. Antibodies specific to RadD (α-RadD) and FomA (α-FomA) are used for detection. FomA serves as a loading control. Molecular weight markers (in kilodaltons) are indicated on the left side of the blot.

Using this strategy, we attempted to generate a new CRISPRi plasmid pZP4C(*radD*), which targets to *radD*. This gene encodes a primary fusobacterial adhesin responsible for *F. nucleatum*’s aggregation with early dental plaque colonizers such as *Actinomyces oris*. It is the final in a four-gene operon (Fig. 2B). To do so, we first identified a 20-nt *radD* target sequence adjacent to a protospacer adjacent motif (PAM) NGG, characteristic of the *S. pyogenes* Cas9 system (Fig. 2B). This sequence was incorporated into the P1 primer’s variable region (Table 2), producing P1(*radD*) (Fig. 2B). This primer, alongside P2 (Fig. 2A & B), was used for PCR amplification of a *radD*-specific sgRNA fragment. It was then inserted into the linearized pZP4C from pZP4C-mCherry via Gibson assembly. Following assembly, we transformed the product into *E. coli* DH5α competent cells. The ChemiDOC MP imaging system was employed under a Cy3 setting to distinguish between colonies. Positive colonies appeared gray, and negatives were stark white (a false color) (Fig. 2C). DNA sequencing of eight randomly selected gray colonies confirmed the efficacy of the *mCherry*-facilitated cloning method (data not shown). All housed the accurate CRISPRi plasmids targeted to *radD*.

To test if the pZP4C(*radD*) can repress *radD* expression, we introduced this plasmid into the wild-type ATCC 23726 strain, resulting in WTpZP4C(*radD*). It was then cultured in a TSPC medium with varied theophylline concentrations. As cultures reached the stationary phase, cells were harvested, washed, and readied for coaggregation tests with *A. oris*. We hypothesized that increasing inducer concentrations would correspondingly impair the aggregation ability. Consistently, cultures treated with 2 mM of theophylline showed a total loss of coaggregation with *A. oris*. Interestingly, even at lower inducer concentrations, aggregation was also completely abolished (Fig. 2D). In contrast, in the absence of the inducer, the cells demonstrated strong aggregation, akin to the wild-type strains (Fig. 2D). We performed Western blotting on the identical cell batches to further substantiate these observations, using a RadD-specific antibody. The results mirrored the coaggregation phenotypes. Cells exposed to inducers exhibited significantly diminished RadD levels, with those at 2 mM inducer showing the most pronounced reduction. The RadD levels in pZP4C(*radD*) cells without inducers were comparable to wild-type strains (Fig. 2E&F). This consolidated evidence further underscores the CRISPRi system’s efficacy in suppressing gene expression in *F. nucleatum*.

### Use of the CRISPRi system for targeting essential gene *bamA*

As our experiments above have shown, the CRISPRi system can effectively inhibit the expression of non-essential genes such as *ftsX* and *radD*. Our next question is whether this system can also suppress essential genes. To address this, we chose *bamA* as the target. This gene encodes the protein BamA, playing a pivotal role in outer membrane protein biogenesis, and its deletion proves lethal for *E. coli* cells (45). In *F. nucleatum*, the *bamA* is the first gene in a quartet, followed by *skp*, *lpxD*, and *glpQ* (Fig. 3A). These genes have diverse roles: *skp* prevents aggregation of outer membrane proteins (46), *lpxD* contributes to lipid A biosynthesis (47), and *glpQ* plays a role in phospholipid degradation (48). Another compelling reason for choosing *bamA* was our prior endeavor to delete this gene from the chromosome. We aimed to discern its role in surface presenting RadD, the fusobacterial adhesin that stands central to our research. We employed our laboratory’s *galK-*based in-frame deletion technique (25). However, all strains we isolated persisted with the wild-type allele and no mutant, suggesting that *bamA* is essential in *F. nucleatum* (not published data).

**Figure 3.**
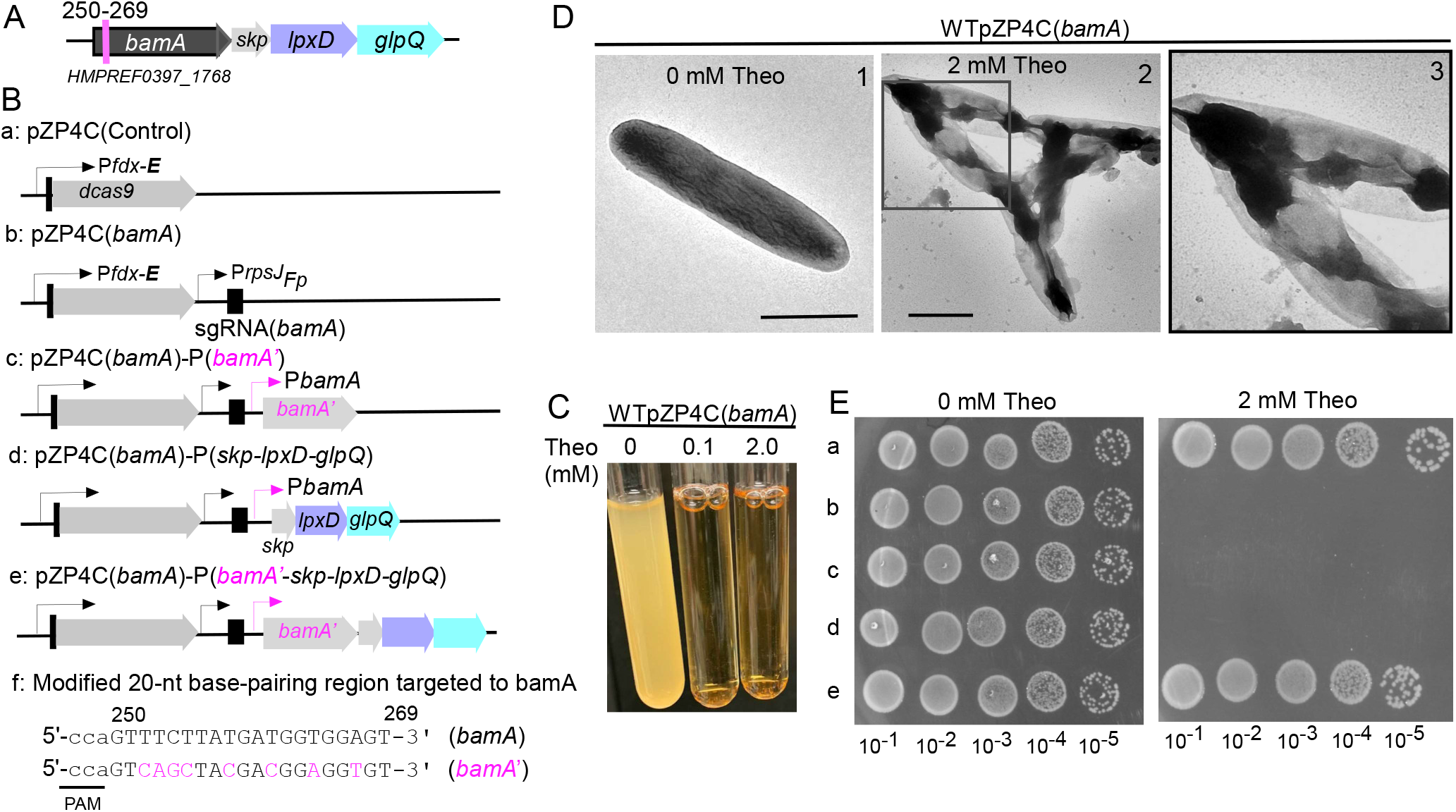
CRISPRi targeting essential gene *bamA*. **(A)** Shown is the *bamA* gene operon in *F. nucleatum*. This operon consists of four consecutive genes: *bamA*, *skp*, *lpxD*, and *glpQ*. The specific 20-nt base-pairing region within the *bamA* gene, which is targeted, is delineated by a red line. The locus number for *bamA* is also provided for reference. **(B)** A schematic diagram detailing the design of various CRISPRi plasmids related to *bamA*. This includes pZP4C(*bamA*) (b), as well as a control plasmid (a) containing *dCas9* without *sgRNA*. Additionally, three modified plasmids derived from pZP4C(*bamA*) are depicted: expressing *bamA* (c), or the triplet genes *skp-lpxD-glpQ* (d), and the entire four gene operon (e). Notably, the sequence expressing *bamA* in c and e has been modified to prevent CRISPRI recognition, with altered nucleotides highlighted in red. The modified *bamA* is named *bamA’*. **(C)** CRISPRi-mediated silencing of *bamA* leads to growth inhibition. Wild-type cells containing pZP4C(*bamA*) were cultivated in a TSPC medium supplemented with varying concentrations (0, 0.1, and 2.0 mM) of theophylline for 22 hours. **(D)** *bamA* silencing induces short filamentation and alters the outer membrane structure. Cells harboring pZP4C(*bamA*) with either 0-or 2-mM theophylline were immobilized on carbon-coated nickel grids, stained with 0.1% uranyl acetate, and observed using a transmission electron microscope. Enlarged views of specific areas (D2) are displayed in D3: scale bar, 1μm. **(E)** The rescue of the death phenotype caused by CRISPRi-mediated *bamA* silencing is demonstrated. Cells containing one of the various plasmids shown in panel (B) were cultured overnight without inducers. Subsequently, these cultures were serially diluted ten-fold and spotted on agar plates with and without inducers. Photographs of the plates were taken after incubation in anaerobic conditions for 3 days.

To test if the CRISPRi system can repress the *bamA* gene expression, we identified a distinct 20-nt sequence within *bamA* for base pairing (Fig. 3A). Using this sequence, we crafted a *bamA*-specific sgRNA and integrated it into the pZP4C-*mCherry* derivative backbone, leading to the generation of the CRISPRi plasmid, pZP4C(*bamA*) (Fig. 3B). Following the transformation of this plasmid into ATCC 23726, we produced the strain WTpZP4C(*bamA*). Remarkably, WTpZP4C(*bamA*) growth trials inhibited entirely growth at a mere 0.1 mM of theophylline (Fig. 3C). To further understand this, we examined its surface structure in the presence of the inducer. Given the growth halt in the inducer’s presence, we increased the inoculation volume when seeding WTpZP4C(*bamA*) to ensure a sufficient quantity of cells for analysis. This way, despite no proliferation, we could still obtain adequate bacterial cells for examination. After an 8-hour growth, we separately harvested the WTpZP4C(*bamA*) cells from both batches – those cultivated without and those with 2 mM theophylline. The harvested cells were stained with uranyl acetate and observed under an electron microscope. In alignment with the established function of *bamA*, the outer membrane of cells grown with theophylline was compromised, rendering the cells more transparent and forming short chains. In contrast, cells without the inducer retained the characteristics of the wild-type (Fig. 3D).

Given the documented polar effect of the CRISPRi systems (49, 50), one may argue that the cell death observed when targeting *bamA* with CRISPRi might stem from the inadvertent silencing of *lpxD* expression, especially since *lpxD* – a downstream gene of *bamA* – is recognized as essential in *E. coli* and other Gram-negative bacteria (51, 52). To explicitly ascertain *bamA’s* essentiality in *F. nucleatum*, we engineered a modified gene version of *bamA* (*bamA’*, Fig 3Bf). This adapted *bamA* underwent modification by introducing several silent mutations to our used 20-nt base-pairing region, allowing it to sidestep CRISPRi targeting (Fig. 3Bf). However, even with this alternation, simultaneous expression of the modified *bamA* (Fig. 3Bc & 3Ec) didn’t prevent cell death in the strain WTpZP4C(*bamA*) when exposed to the inducer, signaling the polar effect of the CRISPRi system. Furthermore, when the other three genes were co-expressed with CRISPRi (Fig. 3Bd & 3Ed), WTpZP4C(*bamA*) ’s growth remained inhibited. Yet, a shift was observed when the entire operon (Fig.3Be) was expressed: WTpZP4C(*bamA*) exhibited normal growth when exposed to theophylline (Fig. 3Ee). This growth pattern was consistent with a control strain (Fig. 3Ba & 3Ea) with a plasmid expressing only *dcas9,* devoid of sgRNA. These data underscore *bamA*’s essentiality and highlight the CRISPRi system’s prowess in effectively silencing essential genes in *F. nucleatum*.

### Use of the CRISPRi system for targeting essential gene *ftsZ*

Exploring essential genes in bacteria like *F. nucleatum* using conventional methodologies has always been fraught with challenges, often tedious and time-intensive (40). In light of this, employing the CRISPRi system to suppress the *bamA* gene marked a significant stride forward. To broaden our understanding of this system’s adaptability, we constructed another plasmid, pZP4C(*ftsZ*) (Fig. 4B), explicitly targeting the *ftsZ* gene. This gene, crucial in almost all bacterial divisions and essential for cell viability in many bacteria (53, 54), is the concluding gene in a three-gene operon in *F. nucleatum* (Fig. 4A). It is preceded by *ftsQ* and *ftsA*, both of which also play important roles in cell division.

**Figure 4.**
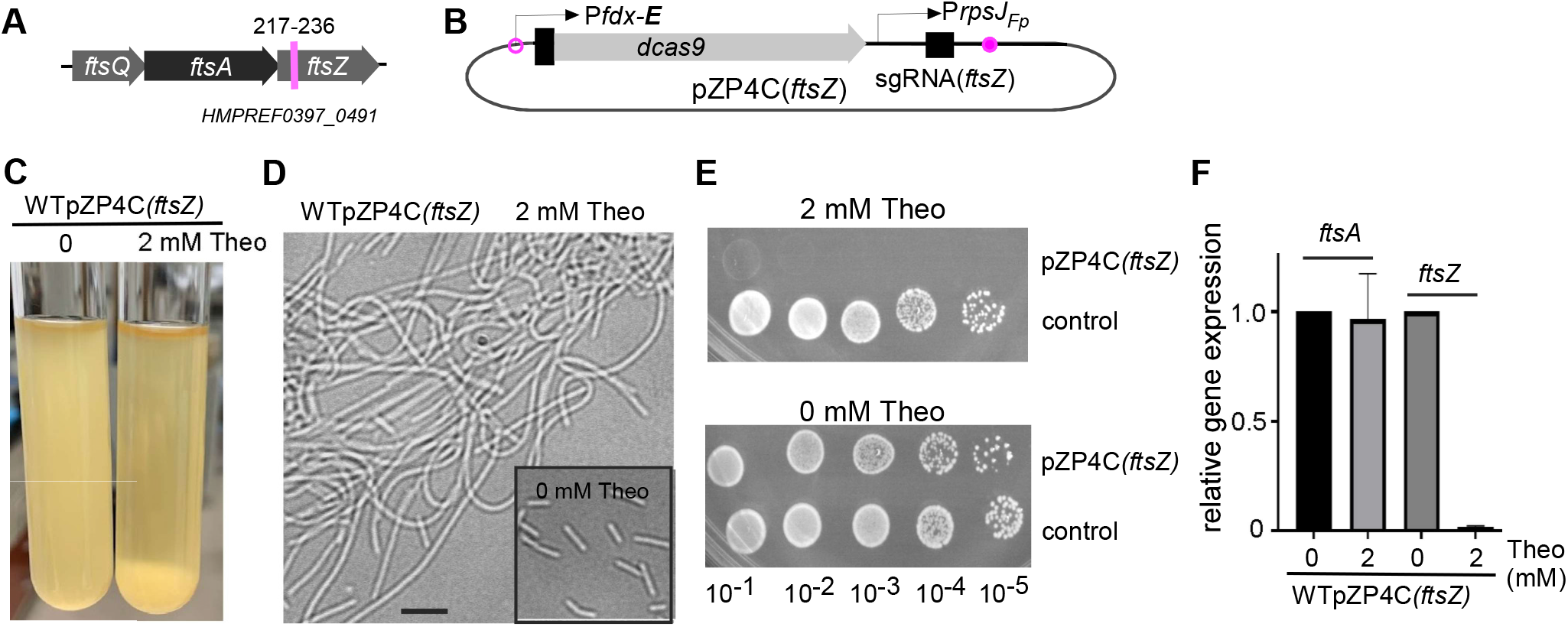
CRISPRi targeting essential gene *ftsZ*. **(A)** The *ftsZ* gene, the last gene of a three-gene operon, is depicted. The specific 20-nt base-pairing region targeted within the *ftsZ* gene is highlighted by a red line. The locus number for *ftsZ* is also provided for reference. **(B)** Display of the pZP4C(*ftsZ*) plasmid design, specifically engineered to target the *ftsZ* gene. **(C)** The aftermath of CRISPRi-mediated *ftsZ* silencing shows most cells sedimented at the bottom of the culture tube. Wild-type cells containing pZP4C(*ftsZ*) were cultivated overnight in a TSPC medium with and without inducers. **(D)** CRISPRi-mediated *ftsZ* silencing causes cell filamentation. Cells from the (C) experiment were observed under a phase-contrast microscope. The insert graph shows cells from cultures grown without inducers as a control. **(E)** Real-time PCR results underline the specific targeting of *ftsZ* by CRISPRi and its minimal polar effect on *ftsA*, the preceding gene. Total RNA, extracted from cells during the early stationary phase with or without inducers, served as the template for the RT reaction. **(F)** The inhibitory growth on agar plates when silencing *ftsZ* using CRISPRi. Wild-type strains with pZP4C(*ftsZ*) or the control plasmid (see Figure 3Ba) were grown in a TSPC medium without inducers. After serial dilutions, samples were spotted on TSPC agar plates with or without 2 mM theophylline. Photographs of the plates were taken after a 3-day incubation period.

We introduced the pZP4C(*ftsZ*) plasmid into the ATCC 23726 strain, yielding the WTpZP4C(*ftsZ*) strain. When cultured with 2 mM theophylline, most of the fusobacterial cells sank to the bottom of the culture. This sedimentation rendered the supernatant noticeably less turbid compared to the culture without the inducer (Fig. 4C). Microscopically, these theophylline-exposed cells exhibit elongated structures, echoing the characteristic function of *ftsZ* (Fig. 4D). However, in its absence, these cells bore a stark resemblance to the wild-type strain (the inert in Fig. 4D). Intriguingly, although the WTpZP4C(*ftsZ*) strain thrived in a liquid medium with theophylline, it faltered on an agar plate with the identical inducer concentration (Fig. 4E).

Given that the CRISPRi system has shown reverse polar effects in certain bacteria – where a targeted gene’s suppression can inadvertently affect its upstream neighbor in the same operon – we sought to investigate this. Through RNA extraction and subsequent quantitative PCR analysis of pZP4C(*ftsZ*) cells grown with and without theophylline, we found a dramatic decline in *ftsZ* mRNA levels upon theophylline exposure. At the same time, the *ftsA* expression remained unaffected (Fig. 4F). These results accentuate the CRISPRi system’s promise in efficiently targeting essential genes in *F. nucleatum*.

### Use of the CRISPRi system to repress gene expression in resistant strain CTI-2 and *F*. *periodonticum*

It can be challenging to manipulate the genes of most fusobacterial strains because they have low transformation efficiencies. This makes it harder to use conventional gene inactivation methods that rely on homologous recombination. However, we have introduced the CRISPRi system as a potential solution for gene inactivation in resistant strains. While these strains may have extremely low transformation abilities, they are not entirely untransformable, which provides an opportunity for genetic intervention. Our CRISPRi system, based on the replicable vector pCWU6, allows for the regulation of target gene expression and a detailed examination of their functions once a successful transformant is achieved. To test this strategy, we chose two fusobacterial strains, CTI-2 and F. periodonticum strain ATCC 33693, to see if it is practical.

The clinical strain CTI-2, isolated from colorectal cancer tissue (27), is categorized under the *F. nucleatum* subsp. *nucleatum* category (28). To determine the CRISPRi system’s efficacy in CTI-2, we constructed a plasmid, pZP4C(*tnaA_CTI-2_*), targeting the *tnaA* gene (Fig. 5A). This gene encodes the enzyme tryptophanase that produces indole from tryptophan. Notably, *tnaA* is a monocistronic gene (Fig. 5A). We selected *tnaA* primarily due to the straightforward detection of indole (40). When indole undergoes a reaction with para-dimethylaminobenzaldenhyde (DMAB) under acidic conditions, it yields a red dye known as rosindole, whose intensity varies based on its concentration, serving as a reliable indicator of indole levels. We introduced 1 µg of pZP4C(*tnaA_CTI-2_*) into CTI-2 using electroporation, only resulting in 5 transformants. In a parallel experiment, we revealed that the same amount of plasmid can produce more than 8,000 transformants in strain ATCC 23726. This stark contrast underscores CTI-2’s exceptionally low transformation efficiency. One of five transformants was named the strain WTpZP4C(*tnaA_CTI-2_*) and further cultivated in varied theophylline concentrations. From each culture that reached the stationary phase, 200 µl was transferred to a 96-well plate. These samples were combined with 100 µl Kovacs’s regent to facilitate indole detection. As the inducer concentration ascended, a decrease in indole production was discernible, with the color transitioning from red to pink (Fig. 5B). Specifically, at 2 mM theophylline, there was a 12-fold reduction in indole synthesi compared with the culture without inducer (Fig. 5C). It is important to note that theophylline did not influence the growth of CTI-2. As expected, in the absence of any inducer, WTpZP4C(*tnaA_CTI-2_*) exhibited negligible differences when compared to the wild-type strain (Fig. 5B). This data suggests that the CRISPRi system can repress gene expression in CTI-2, though the suppression is not absolute.

*F. periodonticum* is one of approximately 13 fusobacterial species and is phylogenetically closest to *F. nucleatum* (9). The type strain for this species, ATCC 33693, was isolated from a human periodontal lesion (55) and has a limited transformation capability. Using a standard fusobacterial transformation protocol with about 1 µg of pCWU6 (a shuttle plasmid for *F. nucleatum*), one can typically obtain around 10 transformants (39).

**Figure 5.**
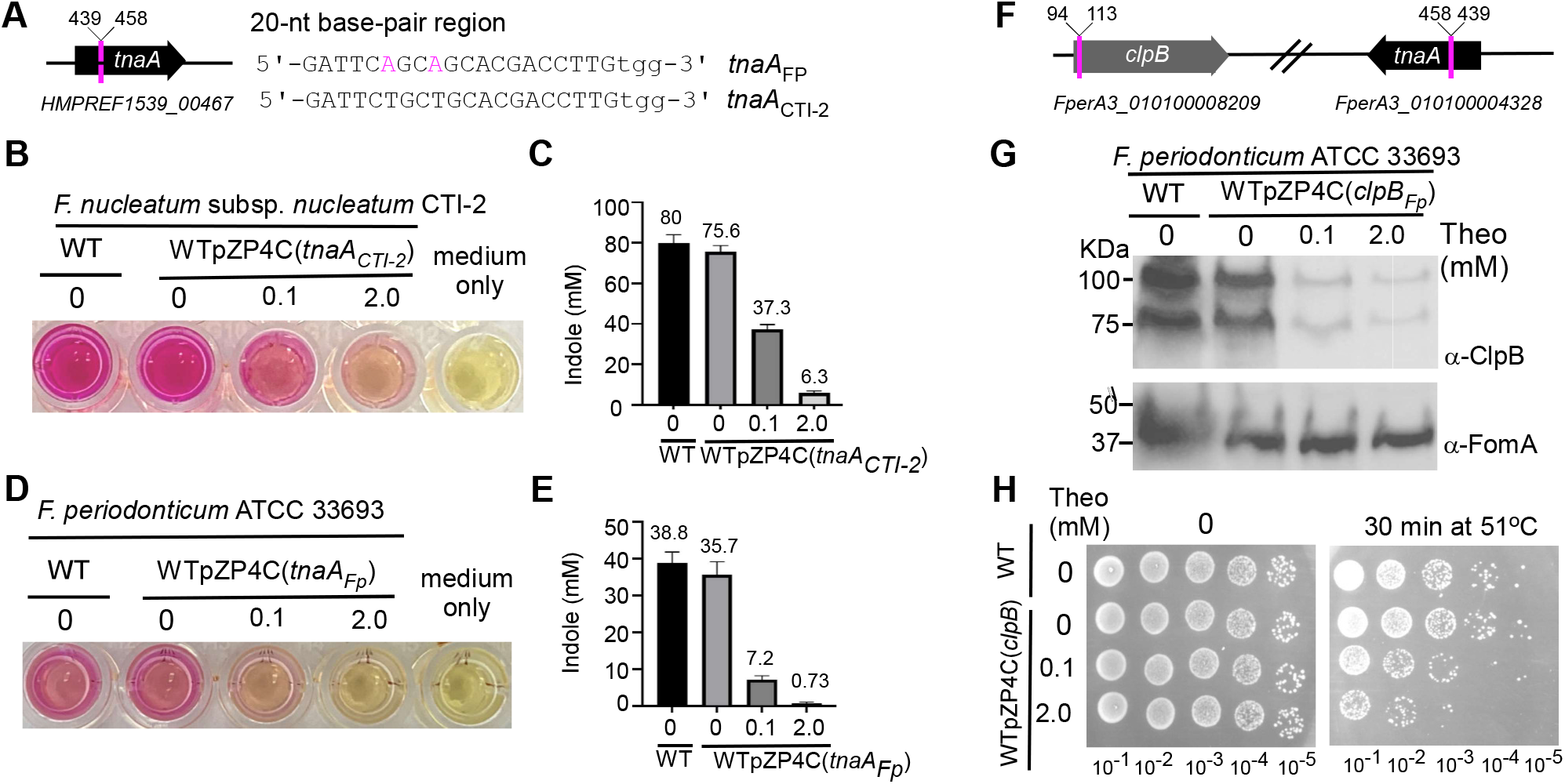
Apply CRISPRi system into *F. nucleatum* clinical strain CTI-2 and *F. periodonticum*. **(A)** The *tnaA* gene locus in the genome of *F. nucleatum* strain CTI-2 is displayed alongside a comparison of the 20-nt base-pairing region targeted within the *tnaA* of CTI-2 with the corresponding region in *F. periodonticum*. **(B)** Diminished indole production due to CRISPRi-mediated *tnaA* knockdown. The strain CTI-2 cells harboring pZP4C(*tnaA_CTI-2_*) are cultivated with varying concentrations of inducers for a 14-hour growth period. Indole production is indicated by introducing Kovac’s reagent, resulting in a red color reaction with indole in the cultures. A WT strain culture is the positive control, while the medium alone is the negative control. **(C and E)** Indole production due to TnaA presence was qualified by indole assay. The measurement of indole production was repeated three times, and the mean values of one representative experiment performed in triplicate are reported and indicated. **(D)** Reduced indole production through CRISPRi-mediated *tnaA* knockdown in *F. periodonticum* wild-type cells harboring pZP4C(*tnaA_Fp_*). As described in (D), an indole assay is employed for validation. **(F)** The *clpB* and *tnaA* loci are illustrated in the genome of *F. periodonticum*, and the location of the targeted 20-nt base pairing regions within the *clpB* and *tnaA* genes are marked by red lines. Corresponding locus numbers for both genes are provided. **(G)** Validation of CRISPRi-mediated *clpB* is confirmed through Western blot analysis. The wild-type cells host pZP4C(*clpB_Fp_*) are cultivated in a TSPC medium supplemented with varying concentrations (0, 0.1, and 2 mM) of theophylline for a 22-hour incubation. Protein samples of equal quantities are separated on a 4-20% Tris-glycine gradient gel and subjected to immunoblotting using anti-ClpB and -FomA antibodies. The latter antibody serves as a loading control. The control group comprised wild-type cells lacking the plasmid. **(H)** Heat sensitivity resulting from CRISPRi-induced *clpB* knockdown is observed. Cells of WTpZP4C(*clpB_Fp_*) are cultured at 37°C with different inducer concentrations overnight. Upon reaching the stationary phase, the cultures are serially diluted in fresh TSPC medium and spotted onto TSPC agar plates lacking inducers. Not-heat-treated cells are included as the control group. A representative experiment of heat sensitivity is shown from three independent replicas.

To test if this CRISPRi system can also manipulate gene expression in *F. periodonticum*, we constructed two plasmids: pZP4C(*tnaA_Fp_*) and pZP4C(*clpB_Fp_*), which target *tnaA* and *clpB*, respectively. ClpB belongs to the heat shock protein 100 family of molecular chaperones and provides resistance to elevated temperature. Both *tnaA* and *clpB* are monocistronic genes (Fig. 5F). It’s worth noting that the 20-nt base-paring region for targeting *tnaA* in *F. periodonticum* differs by only two nucleotides compared to its CTI-2 counterpart (Fig. 5A). Following the transformation of these two plasmids into ATCC 33693, we obtained two distinct strains: WTpZP4C(*tnaA_Fp_*) and WTpZP4C(*clpB_Fp_*). Our initial experiments examined the indole synthesis by WTpZP4C(*tnaA_Fp_*) under three different theophylline concentrations. While *F. periodonticum* naturally produces less indole than CTI-2 (Fig. 5C & 5E), an increasing theophylline concentration led to subsequent decreases in indole production (Fig. 5D). Notably, in the presence of a 2 mM theophylline concentration, the dye’s visual hue closely resembled that of the standard medium (Fig. 5D). Quantitative analysis for indole showed that its production was reduced 50-fold when compared to the culture without the inducer (Fig. 5E).

Next, we tested the ability of the CRISPRi system to repress *clpB* expression in *F. periodonticum*. After cultivating WTpZP4C(*clpB_Fp_*) with various theophylline amounts and harvesting the cells at the stationary phase, Western blot analysis revealed two bands similar to patterns in *E. coli* (56). These represent both the complete and a naturally shortened form of ClpB. A rise in theophylline resulted in a noticeable reduction in ClpB expression, almost vanishing at 2 mM (Fig. 5G), making the cells more sensitive to heat (Fig. 5H). Our data demonstrated the CRISPRi system’s potential in targeting gene repression in CTI-2 and *F. periodonticum* strains.

## DISCUSSION

Gene inactivation in *F. nucleatum* is challenging, and most fusobacterial strains are recalcitrant to DNA manipulation. This study introduces a theophylline-inducible CRISPRi system to address these problems. We demonstrated that this system could effectively repress selected genes’ expression, including nonessential genes (*ftsX* and *radD*) and essential genes (*bamA* and *ftsZ*) in the model organism, ATCC 23726. Most importantly, this CRISPRi approach has allowed us to study gene functions in strains such as CTI-2 and *F. periodonticum*, which are historically resistant to genetic manipulation.

The traditional methods for gene inactivation, such as creating in-frame deletion mutants and insertion mutations, rely on homologous recombination. However, this poses a challenge for all fusobacterial strains (23, 26), which typically have notably poor rates of homologous recombination. To provide some perspective, out of 2,000 plasmids introduced for gene inactivation, only one tends to integrate into the host genome through homologous sequences (39). Thus, the host would ideally need an exceptional plasmid transformation ability for successful integration. Regrettably, the transformation capability of many fusobacterial stains is hampered primarily due to their sophisticated restriction-modification systems (RMs) (23). When *E. coli*-derived plasmids are introduced into these strains via electroporation, they frequently succumb to degradation by these RMs (57). This results in a starkly low transformation rate among many fusobacterial strains. Consequently, many struggle to align with prevailing gene inactivation methods, particularly those tailored for the model organism, ATCC 23726, which boasts impressive transformation efficiency (26).

There are two classical methods for gene inactivation in bacteria with low transformation efficiencies. The first involves in vitro methylation of plasmids using DNA methyltransferase before introduction into the host, protecting the DNA from the host’s restriction enzymes (58, 59). While this method proved effective in the strain ATCC 25586, it’s labor-intensive, and its applicability can be strain-specific (23). The second employs conditional suicide vectors with temperature-sensitive replication origins (60). These vectors replicate at specific temperatures and can integrate into the chromosome at others. However, a tailored plasmid for *F. nucleatum* remains undeveloped.

In recent years, shuttle plasmids with CRISPR systems also become valuable when working with bacterial species that are challenging to manipulate genetically (61, 62). The use of the CRISPR-Cas system for bacterial gene deletion involves designing a shuttle plasmid with CRISPR components and sequences flanking the target gene. Once introduced into the bacteria, the CRISPR system induces a break at the target site, which the cell repairs using the plasmid’s flanking sequences, effectively deleting the gene. We are in the process of adapting this approach for *F. nucleatum* and will present our results in due course. While the methods above modify the gene’s structure, CRISPRi remains an exception. Instead of altering the gene’s structure or relying on homologous recombination, CRISPRi inhibits gene transcription (63). This makes a CRISPRi-based shuttle plasmid viable for fusobacterial strains when the in-frame deletion technique for ATCC 23726 is inaccessible. Despite the transformation challenges in many fusobacterial strains, achieving a single transformant with a shuttle plasmid remains plausible. Our CRISPRi system is built in the shuttle plasmid pCWU6. Our hands-on experiment targeted the *tnaA* gene in the clinical strains CTI-2 and *F. periodonticum* ATCC 33693, resulting in diminished indole levels aligned with inducer concentrations (Fig. 5B& 5D). Interestingly, the suppression efficiency of *tnaA* in CTI-2 was less robust than in ATCC 33693 (Fig 5C & 5E), possibly due to the suboptimal activity of the *rpsJ* promoter from *F. periodonticum* within CTI-2. Thus, for optimal CRISPRi performance, employing the host’s endogenous promoter for sgRNA expression is recommended. The amounts of sgRNA expression affect the suppression effect of CRISPRi (63). Targeting the *clpB* gene in ATCC 33693 led to near-complete silencing under a 2 mM inducer (Fig. 5G), enhancing the heat sensitivity of these cells compared to those without the inducer (Fig. 5H). This research marks a pioneering effort in probing gene functions in CTI-2 and *F. periodonticum*, paving the way for future studies in other challenging strains, notably those in subsp. *animals*. Given the prevalence of subsp. *animals* in the colon and its frequent association with colorectal cancer, and the genetic intractability of this bacterial group, there is a pressing need to identify a representative strain for subsp. *animals*, especially as ATCC 23726-a non-colon inhabitant-is currently the model for colon cancer pathogenesis.

The foremost advantage of CRISPRi is its simplicity. One only needs to substitute the sgRNA to target a new gene, a task easily achieved through PCR. Moreover, including a mCherry reporter in the CRISPRi cloning plasmid simplifies the process of screening for *E. coli* containing the correct (positive) plasmids (Fig. 2A). Thus, CRISPRi is particularly well-suited for high-throughput studies of gene functions on a whole genome scale. While the Tn5-based transposon also offers a method for genome-scale gene function studies, it falls short in investigating essential genes due to insertional lethality (25). In contrast, CRISPRi provides a targeted approach, enabling the analysis of essential genes by suppressing their expression without altering the DNA sequence (64). Essential genes in bacteria are vital for its survival and proper functioning under standard conditions. Their disruption often results in lethality or impaired growth. Due to their central roles in bacterial physiology, essential genes are often explored as potential antibiotic targets. Recently, we deleted *lepB*, an essential gene in *F. nucleatum* (40). This deletion from the chromosome was made possible by introducing an extra functional *lepB* copy via an expression vector. While this method was effective, it was also lengthy, taking us nearly a month to produce the deletion strain. On the other hand, using CRISPRi to study an essential gene is more time-efficient, demanding just a simple PCR to modify the sgRNA.

The BAM complex in Gram-negative bacteria, integral for protein folding and insertion into the outer membrane, consists of BamA and its BamB-E accessory proteins in *E. coli* (65) and is essential for bacterial survival (45), positioning it as a potential antibacterial therapy target. Intriguingly, *F. nucleatum* only possesses the *bamA* gene, lacking the associated accessory protein homologs (https://img.jgi.doe.gov/). When *bamA* is targeted with CRISPRi, cells show no growth in the presence of the inducer (Fig. 3C). Electron microscopy reveals a significant compromise in outer membrane integrity, hindering effective cell division (Fig. 3D). The *ftsZ* gene in bacteria encodes the FtsZ protein, which forms a Z-ring at the cell’s midpoint, playing a crucial role in bacterial cell division (66). Using CRISPRi to target *ftsZ* causes cells to elongate in broth cultures (Fig. 4D). It prevents growth on agar plates (Fig. 4F), mirroring observations made with *E. coli* FtsZ, where its depletion similarly results in extended cell structures and halted growth on solid media (53). The CRISPRi system is instrumental in probing essential genes in *F. nucleatum*. With the pCWU6 vector’s stability (25, 26) and the FDA-approved status and well-tolerated nature of theophylline in mice and guineas pigs, our theophylline-responsive riboswitch-controlled CRISPRi, especially when targeting essential genes, holds promise for animal studies exploring fusobacterial pathogenesis. This system could also pave the way for developing conditional live attenuated vaccines, targeting essential genes like *bamA* or *clpB*, where the bacteria can be selectively killed using the inducer after eliciting an immune response. Given the simplicity and effectiveness of the CRISPRi system, our lab is currently developing a CRISPRi library targeting essential genes in the model organism ATCC 23726. A comprehensive CRISPRi-driven analysis of these genes in *F. nucleatum* is poised to provide deeper insights into its biology and pathogenicity.

The CRISPRi system offers distinct advantages but has limitations concerning gene polarity. It struggles when aimed at multi-gene operons’ initial or central genes (64). For example, downstream genes also exhibited inhibited expression when targeting *bamA*, the lead gene in a quartet operon (Fig. 3E). As a result, introducing a *bamA* variant with a 20-nt base paring failed to prevent cell death (Fig. 3Ec), highlighting areas where CRISRPi is less effective than in-frame deletion. However, this polar effect can be mitigated by co-expressing the entire gene operon, minus the targeted gene, within the same plasmid (Fig. 3Ed & 3Ee). While some bacteria show reverse polarity effects with CRISPRi (64), this isn’t evident in *F. nucleatum*. Specifically, when targeting *ftsZ*, the final gene in a triad, preceding gene expression remained unchanged (Fig. 4E). Yet, this finding needs broader validation to determine the full scope of CRISPRi’s effects on *F. nucleatum*’s gene polarity.

In *F. nucleatum*, in-frame deletion mutants for nonessential genes can be produced using either *galK*-based (25, 32) or toxin-based (33, 39) counterselection methods. However, these processes can be time-intensive. The CRISPRi system offers a rapid preliminary evaluation of a gene’s function. Once its role is ascertained, one can invest time in crafting a precise in-frame deletion for a more detailed study.

In summary, the riboswitch-controlled CRISPRi system offers a streamlined approach to investigating genes within *F. nucleatum*, particularly emphasizing essential gene analysis. Moreover, our shuttle plasmid-based CRISPRi design circumvents transformation barriers, enabling gene activation in previously recalcitrant strains of *F. nucleatum* and expanding its applicability to other fusobacterial species. This advances our understanding of fusobacterial genetics and biology to the subspecies and even strain-specific level, positioning us closer to devising innovative strategies against these notoriously opportunistic pathogens.

## MATERIALS AND METHODS

### Bacterial strains, plasmids, and growth conditions

A comprehensive list of bacterial strains employed in this study is outlined in Table 1. Culturing both *F. nucleatum* and *F. periodonticum* strains was carried out using a TSPC medium comprising 3% tryptic soy broth (TSB, BD) and 1% Bacto^TM^ peptone. Before the inoculation, a 0.05% cysteine supplement was meticulously added to the TSPC broth medium or agar plates. The cultures were propagated within an anaerobic chamber filled with a gas mixture of 80% N_2_, 10% H_2,_ and 10% CO_2_. *Escherichia coli* strains were grown in Luria-Bertani (LB) broth. The fusobacterial strains harboring CRISPRi plasmids were cultivated in the presence of thiamphenicol (5 µg/ml) while being exposed to various concentrations of inducer (theophylline). A solution of 40 mM theophylline was prepared in a TSP medium, stored at 4°C for long-term storage, and prewarmed to 37°C before use. When required, Antibiotics used as needed were chloramphenicol (15 µg ml^-1^) and thiamphenicol (5 µg ml^-1^). Reagents were purchased from Sigma unless indicated otherwise.

**Table 1:** Bacterial strains and plasmids used.

| Strain & Plasmid | Description | Reference |
| --- | --- | --- |
| Strains |  |  |
| *F. nucleatum* 23726 | Parental strain (wild-type strain) | From ATCC |
| *F. nucleatum* CTI-2 | Wild-type strain, isolated from human CRC samples | (27) |
| *F. periodonticum* 33693 | Wild-type strain | From ATCC |
| *F. nucleatum* CW2 | $\Delta ftsX$; an isogenic derivative of CW1 | (25) |
| *F. nucleatum* BG01 | $\Delta radD$; an isogenic derivative of 23726 | (39) |
| *F. nucleatum* ZP05 | WT strain 23726 containing pZP4C | This study |
| *F. nucleatum* ZP06 | WT strain 23726 containing pZP4C(*radD*) | This study |
| *F. nucleatum* ZP07 | WT strain 23726 containing pZP4C(*bamA*) | This study |
| *F. nucleatum* ZP07a | WT strain 23726 containing pZP4C(*control*) | This study |
| *F. nucleatum* ZP07b | WT strain 23726 containing pZP4C(*bamA*)-P-*bamA*' | This study |
| *F. nucleatum* ZP07c | WT strain 23726 containing pZP4C(*bamA*)-P-*skp-lpxD-glpQ* | This study |
| *F. nucleatum* ZP07d | WT strain 23726 containing pZP4C(*bamA*)-P-*bamA*'-*skp-lpxD-glpQ* | This study |
| *F. nucleatum* ZP08 | WT strain 23726 containing pZP4C(*ftsZ*) |  |
| *F. nucleatum* ZP09 | WT strain 33693 containing pZP4C(*clpB<sub>FP</sub>*) | This study |
| *F. nucleatum* ZP10 | WT strain 33693 containing pZP4C(*tnaA<sub>FP</sub>*) | This study |
| *F. nucleatum* ZP11 | WT strain CTI-2 containing pZP4C(*tnaA<sub>CTI-2</sub>*) | This study |
| *E. coli* DH5α | cloning host | From NEB |
| Plasmids |  |  |
| pCWU6 | *E. coli*/*Fusobacterium* shuttle vector, chloramphenicol /thiamphenicol resistance; *cm<sup>R</sup>/thia<sup>R</sup>* | (25) |
| pRECas1 | The *fdx* promoter incorporating the riboswitch element E controls Cas9 expression. | (44) |
| pIA33 | *Pxyl::dcas9-opt Pgdh::sgRNA-rfp catP* | (67) |
| pZP4C | *Pfdx::dcas9-opt PrpsJ<sub>FP</sub>::sgRNA-ftsX catP* | This study |
| pZP4C(*radD*) | *Pfdx::dcas9-opt PrpsJ<sub>FP</sub>::sgRNA-radD catP* | This study |
| pZP4C-*mCherry* | *Pfdx::dcas9-opt PrpsJ<sub>FP</sub>::PrpsJ<sub>cd</sub>::mCherry catP* | This study |
| pZP4C(*control*) | *Pfdx::dcas9-opt catP* | This study |
| pZP4C(*bamA*) | *Pfdx::dcas9-opt PrpsJ<sub>FP</sub>::sgRNA-bamA catP* | This study |
| pZP4C(*bamA*)-P-*bamA*' | *Pfdx::dcas9-opt PrpsJ<sub>FP</sub>::sgRNA-bamA PbamA-bamA' catP* | This Study |
| pZP4C(*bamA*)-P-*skp-lpxD-glpQ* | *Pfdx::dcas9-opt PrpsJ<sub>FP</sub>::sgRNA-bamA PbamA-skp-lpxD-glpQ catP* | This study |
| pZP4C(*bamA*)-P-*bamA*'-*skp-lpxD-glpQ* | *Pfdx::dcas9-opt PrpsJ<sub>FP</sub>::sgRNA-bamA PbamA-bamA-skp-lpxD-glpQ catP* | This study |
| pZP4C(*ftsZ*) | *Pfdx::dcas9-opt PrpsJ<sub>FP</sub>::sgRNA-ftsZ catP* | This study |
| pZP4C(*clpB<sub>FP</sub>*) | *Pfdx::dcas9-opt PrpsJ<sub>FP</sub>::sgRNA-clpB<sub>FP</sub> catP* | This study |
| pZP4C(*tnaA<sub>FP</sub>*) | *Pfdx::dcas9-opt PrpsJ<sub>FP</sub>::sgRNA-tnaA<sub>FP</sub> catP* | This study |
| pZP4C(*tnaA<sub>CTI-2</sub>*) | *Pfdx::dcas9-opt PrpsJ<sub>FP</sub>::sgRNA-tnaA<sub>CTI-2</sub> catP* | This study |

### Plasmid and strain construction

All constructed plasmids in this study are listed in Table 1. These plasmids were methodically created utilizing the Gibson assembly technique with the NEBuilder^®^ HiFi DNA Assembly Master Mix from New England Biolabs, as per the manufacturer’s guidelines. In brief, a linearized cloning vector (ranging from 50 to 100 ng), derived from enzymatic digestion by two restriction enzymes (SacI/HindIII or MscI/NotI), was combined with 1-4 PCR cloning fragments (approximately 120 ng each). This cloning vector/PCR fragment(s) mixture was then added to 10 µl of 2x Gibson Assembly Master Mix, achieving a total volume of 20 µl. The Gibson assembly process was performed at 50°C for 20-60 minutes. Following this, 5 µl of the assembled product was used to transform *E. coli* DH5α competent cells. The authenticity of the resultant plasmids was verified through DNA sequencing. Subsequently, electroporation transferred these validated plasmids to fusobacterial strains (38). The oligonucleotide primers listed in Table 2 were custom-synthesized by Sigma Aldrich.

i. pZP4C. Two primary components are essential in constructing an efficient CRISPRi plasmid: *dcas9* and *sgRNA*. Typically, the *dcas9* is designed for expression under an inducible promoter system, while *sgRNA* expression operates under a constitutively active promoter. We planned to use a riboswitch-based inducible system to modulate *dcas9* expression and harness the *rpsJ* promoter from *F. periodonticum* – a robust and active promoter – to steer *sgRNA* production. To this end, we designed four primer pairs: 1Pfdx-E-F/2Pfdx-E-R, 3dcas9-F/4dcas9-R, rpsJfp-F/R, gDNA(ftsX)-F/gDNA-R. Using these primers, we amplified the fragments Pfdx-E, dcas9, PrpsJFp, and sgRNA. For the PCR reaction, the DNA templates employed were plasmid pREcas1(44), plasmid pIA33 (67), the genomic DNA of *F. periodonticum*, and pIA33, in order. Post-PCR, all four fragments underwent gel purification. Approximately 120 ng from each fragment was pooled, then mixed with 50 ng of the SacI/HindIII-digested pCWU6 – an *E. coli*/*F. nucleatum* shuttle vector integrated with a chloramphenicol resistance marker. This combined solution was treated with an equal volume of 2x Gibson assembly master mix. The assembled construct was later introduced into *E. coli* DH5α cells using standard transformation procedures. The newly generated plasmid was designed as pZP4C. DNA sequencing verified its sequence. Given that the primer gDNA(*ftsX*)-F contains a distinct 20-nt sequence aimed at *ftsX*, we projected that pZP4C would silence fusobacterial *ftsX* gene expression upon adding inducers to the culture medium.
ii. pZP4C-mCherry. To facilitate efficient Gibson assembly and streamline the screening of positive clones, we developed the plasmid of pZP4C-*mCherry*. This plasmid carries the *mCherry* gene, which encodes a red fluorescent protein. Within the plasmid, *mCherry* is positioned at the *sgRNA* site, bookended by MscI/NotI restriction sites on each end. This configuration allows for easy cloning of *sgRNAs* and can be swapped out to target various genes. We began by linearizing the pZP4C plasmid to create this construct using MscI/NotI enzymes. Simultaneously, the *mCherry* gene-driven by an *rpsJ_Cd_* promoter – was amplified from the pCWU6 (25) template using the mCherry-F/R primer pair. Each primer in this pair was designed with a 22-bp overlapping sequence at the 5’. This overlap is complementary to the terminal arms of the pZP4C backbone, and its purpose is specifically to aid the Gibson assembly process. For the Gibson assembly, 120 ng of *mCherry* amplicon was combined with 50 ng of the aforementioned linearized pZP4C. Bacteria transformed with the resultant pZP4C-*mCherry* produce red colonies, a phenotype attributed to mCherry expression.
iii. pZP4C(*radD*), pZP4C(*bamA*), pZP4C(*ftsZ*), pZP4C(*clpB_Fp_*), pZP4C(*tnaA_Fp_*) and pZP4C(*tnaA_CTI-2_*). To create CRISPRi plasmids, we substituted the *mCherry* gene in the PZP4C-*mCherry* vector with the targeted *sgRNA*. This *sgRNA* can be amplified via PCR using a gene-specific forward primer P1 and a universal reverse primer P2. Primer P1 is composed of three distinct segments (5’ to 3’): A 20-nt sequence in the *rpsJ* promoter end, aiding in Gibson assembly, a 20-nt base-paring sequence specific to the target gene, and a 19-nt sequence from sgRNA handle region. The design for P1 is as follows: GTAGTTAATTTAAAATGGCCAxxxxxxxxxxxxxxxxxxxxGTTTTAGAGCTAGAAATAG. The 20-“X” nucleotide denotes the 20-nt targeted gene-specific region. When shifting targets, only this 20-nt sequence requires alternation. To identify suitable 20-nt base-pairing sequences for the targets, The Eukaryotic Pathogen CRISPR guide RNA/DNA Design tool ((http://grna.ctegd.uga.edu/) was employed. Even though this tool was initially developed for eukaryotic pathogens, it was equally adept for bacterial pathogens. Our reference was the Fungi (*A. aculeatus* ATCC 16872) genome. Using standard parameters, we located 20-nt guides that concluded with the NGG Protospacer Adjacent Motif (PAM) at their 3’ terminus in the template strand. Selection criteria for the guides included high CRISPRater efficiency scores, optimal transcription conditions, and binding to the coding strand in the initial third of the gene. To prevent possible off-target effects, the candidate sequence was evaluated using NCBI-Blast to ensure no matches in the PAM-proximal region, as dCas9 is intolerant to mutations there (49). We embedded each of the identified 20-nt sequences for these target genes into the P1 primer for the PCR step. Employing pZP4C as our template and teaming it with P2, we amplified the specific sgRNA for each target gene. Using Gibson assembly, the derived 196-bp PCR product was cloned into the MscI/NotI-digested pZP4C-mCherry, effectively replacing the *mCherry* gene and birthing the CRISPRi plasmid. It is worth noting that within the *mCherry* gene, another MscI site is present. Hence, digesting pZP4C-*mCherry* with MscI/NotI results in two distinct fragments (approximately 500bp and 700bp) in addition to the main plasmid backbone. *E. coli* cells hosting the new CRISPRi plasmids manifested as gray colonies. However, colonies stemming from either non-digested or self-connected pZP4C-*mCherry* radiated a sharp white when viewed under the Cy3 setting of the ChemiDOC^TM^ MP Imaging System by Bio-Rad, a result of the *mCherry* expression. Finally, to ensure accurate *sgRNA* incorporation, all produced CRISPRi plasmids underwent sequencing validation.
iv. pZP4C(control). To construct a CRISPRi control plasmid containing only *dcas9* and lacking *sgRNA*, we employed the primer pair 1Pfdx-E-F/dcas9-R2 to amplify *dcas9* alongside its promoter system *Pfdx-E* from pZP4C. The produced *Pfdx-E-dcas9* fragment was subsequently cloned into the SacI/HindIII-cut pCWU6, yielding the resultant plasmid, pZP4C(control).
v. pZP4C(*bamA*)-P-*bamA’*, pZP4C(*bamA*)-P-*skp-lpxD-glqQ*, and pZP(*bamA*)-P-*bamA’-skp-lpxD-glqQ*. In constructing pZP4C(*bamA*)-P-*bamA’*, we aimed to express the *bamA* gene without interference from the CRISPRi mechanism. To this end, silent mutations were introduced within the 20-nt region of the *bamA* gene targeted by CRISPRi. The *Pfdx-E-dcas9-PrpsJFn-sgRNA* (CRISPRi unit) was amplified from pZP4C(*bamA*) using primers 1Pfdx-E-F and gRNA-R2. A segment from the *bamA* gene’s native promoter to the targeted 20-nt region (segment 1) was PCR-amplified using primers bamA-up-F/R. The 5’ end of bamA-up-R was engineered to include the silent mutations of the 20-nt sequence. A subsequent segment of the *bamA* gene, spanning from the mutated 20-nt region to the gene’s termination (segment 2), was amplified using primers bamA-dn-F/R, with the 5’ end of bamA-dn-F similarly incorporating the silent mutations. Both segments of *bamA* contained the mutated 20-nt sequence, serving as an overlap for Gibson assembly. The three PCR products were cloned into SacI/HindIII-digested pCWU6 through Gibson assembly, resulting in pZP4C(*bamA*)-P-*bamA’*. To generate pZP4C(*bamA*)-P-*skp-lpxD-glqQ,* the *bamA* promoter region was amplified using primers bamA-up-F/PbamA-upR. A PCR product encompassing gene *skp*, *lpxQ*, and *glqQ* was procured using primers skp-F/glpQ-R. These fragments and the CRISPRi unit were cloned into SacI/HindIII-cut pCWU6 to produce the desired plasmid. To make pZP(*bamA*)-P-*bamA’-skp-lpxD-glqQ*, a region from segment 2 to *glqQ* was amplified using primers bamA-dn-F and glqQ-R. This PCR fragment, segment 1, and the CRISPRi unit were cloned into pCWU6 through Gibson assembly to finalize the construct.

**Table 2:**
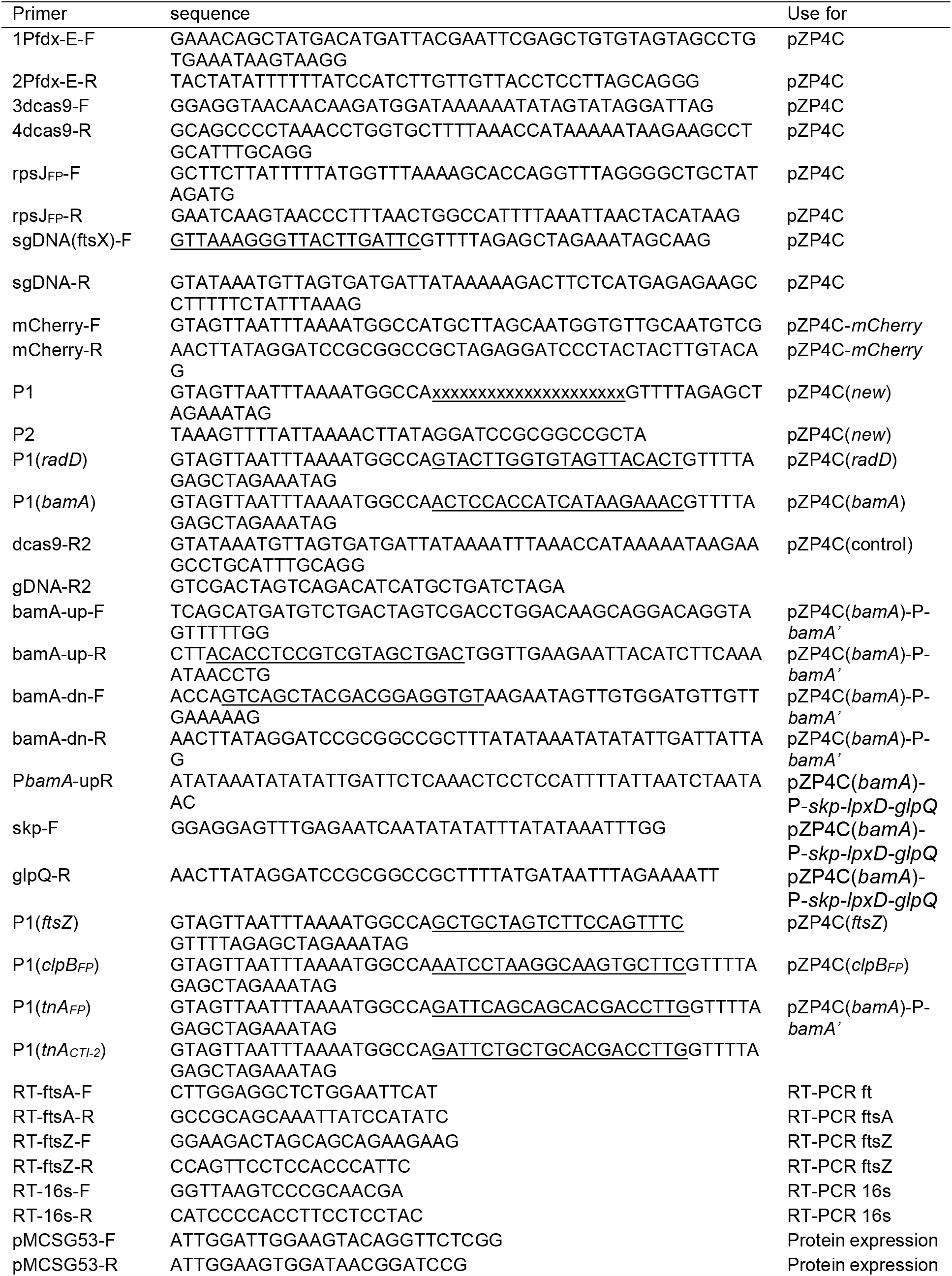
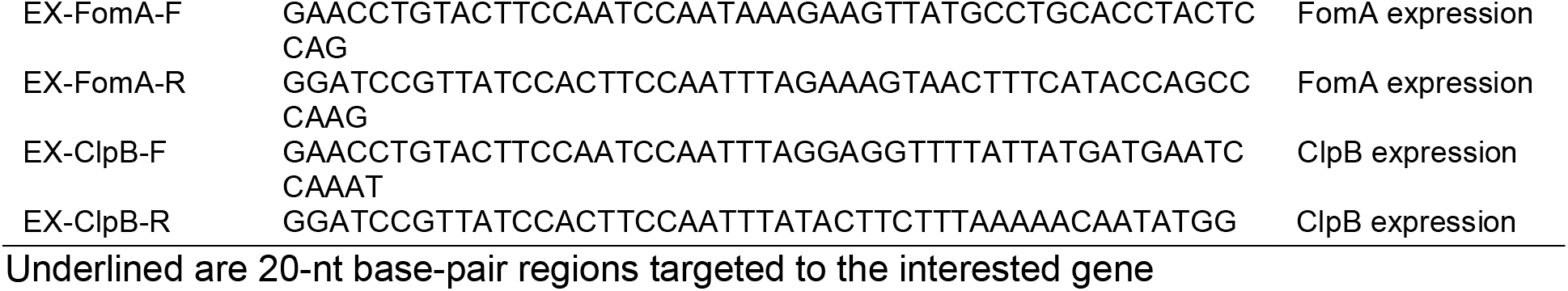
Primers used in this study.

All generated plasmids were authenticated by sequencing and subsequently introduced into *F. nucleatum* ATCC 23726, CTI-2 strain, or *F*. *periodonticum* ATCC 33693, following the electroporation protocol we discussed in a previous publication (38). In brief, cells from a 100 ml stationary phase culture of either ATCC 23726, CTI-2, or *F. periodonticum* were collected via centrifugation. After being washed twice with sterile water and one with 10% glycerol, they were suspended in 3ml of 10% glycerol and aliquoted into 0.2 ml samples. These samples were rapidly frozen with dry ice and preserved at −80°C. We added 1.0 mg of various CRISPRi plasmids to the 0.2 ml fusobacterial competent cells in a chilled cuvette (0.1 cm electrode gap, Bio-Rad), allowing it to rest on ice for 10 minutes. Electroporation was performed using the Bio-Rad Gene Pulser II, adjusted to 25 kV/cm, 25 F, and 200 Ω. Immediately after electroporation, the cells were extended in 1 ml of pre-reduced, pre-warmed TSPC medium. After an anaerobic incubation for 5 hours, the culture was spread on TSPC agar plates containing 5 µg/ml thiamphenicol.

### Theophylline-inducible assay

All fusobacterial strains containing the CRISPRi plasmid were initially cultured in a TSPC medium supplemented with 5 µg/ml thiamphenicol, without any inducer, and allowed to grow overnight. The overnight culture was then diluted at a 1:1000 ratio and incubated in a fresh TSPC medium with the desired concentration of theophylline for 22 hours. Alternatively, cultures were 10-fold serially diluted, and 7 µl from each dilution was spotted onto plates with or without 2 mM theophylline. Following spotting, the plates were incubated for 3 days in an anaerobic chamber before being photographed.

### Western blotting analysis

For FtsX detection in *F. nucleatum*, 22-hour cultures of strains containing pZP4C at various theophylline concentrations (0, 0.1, and 2 mM) were vortexed to evenly distribute the cells, countering the filamentation and tangling observed from *ftsX* silencing via CRISPRi. Post-resuspension, cells from 1 ml of each culture were collected, washed twice with water, and resuspended in the SDS sample buffer. Following a 10-minute boiling, they were subjected to 4-20% Tris-glycine gradient SDS-PAGE and immunoblotting using antibodies against FtsX and FomA (control protein). To generate polyclonal antibodies against FomA, the *fomA* coding region, excluding the signal peptide, was PCR-amplified using primers EX-FomA-F/R. Concurrently, the backbone of the pMCSG53 expression vector (68) was amplified with primes pMCSG53-F/R. Both PCR products were fused via Gibson assembly, resulting in a recombinant plasmid introduced into *E. coli* BL21(DE3). The expressed H6-FomA protein was affinity-purified and utilized for antibody production at Cocalico Biologicals, Inc.

For RadD detection in *F. nucleatum*, cells from strains WT, Δ*radD*, and WT containing pZP4C(*radD*) at varying theophylline concentrations were harvested after a 22-hour growth. These samples were heated to 70°C for 10 minutes, analyzed through 4-20% Tris-glycine gradient SDS-PAGE, and Immunoblotted using previously generated antibodies against RadD (24).

For ClpB detection in *F. periodonticum*, we created a pMCSG53-based *ClpB* expression vector. Using primers EX-ClpB-E/F, the *clpB* gene from *F. nucleatum* ATCC 23726 (HIMPREF0397_1800, available at http://img.jgi.doe.gov/) was amplified and cloned into pMCSG53. This vector was then transformed into *E. coli* BL21(DE3), and the H6-ClpB protein was isolated through affinity chromatography. This purified protein was used to produce antibodies through Cocalico Biologicals, Inc. WT pZP4C(*clpB_Fp_*) cells, cultivated in varying theophylline concentrations, were processed and subjected to SDS-PAGE and immunoblotted using rabbit anti-ClpB and anti-FomA antibodies at 1:1000 dilution.

### Bacterial co-aggregation

Co-aggregation assays were conducted using *F. nucleatum* wild-type, Δ*radD,* or wild-type cells containing pZP4C(*radD*), combined with *A. oris* MG-1, following a previously described method (25). In brief, stationary-phase cultures of bacterial strains were grown in TSPC with/ without inducers or heart infusion broth for MG-1. Post centrifugation, cells were washed and resuspended in coaggregation buffer (200 mM Tris-HCl, pH 7.4, 150 mM NaCl, 0.1 mM CaCl_2_) (69), ensuring an equal cell density of approximately 2 x 10^9^ ml^-1^ based upon OD_600_ values. For the assay, 0.20 ml aliquots of *Actinomyces* and various fusobacterial cell suspensions were mixed in a 24-well plate, briefly shaken on a rotator, and then imaged.

### Electron microscopy

Silencing *bamA* leads to cell death, making it challenging to obtain sufficient *bamA*-depleted cells from electron microscopy, especially when using high dilution ratios like 1:1000. To circumvent this and ensure we gather ample cells for analysis, we first grew WT cells containing pZP4C(*bamA*) in TSPC medium without an inducer overnight. We employed a 1:5 dilution strategy using this overnight culture, expanding the culture in 8ml of TSPC medium. Although *bamA* silencing inhibits cell growth, the generous starting inoculum from the overnight culture compensated for this, ensuring a substantial yield. The culture was subsequently incubated at 37°C in an anaerobic chamber for 8 hours. Following incubation, we centrifuged the culture to collect the cells, washed the resultant pellet, and resuspended it in 0.1 M NaCl. A 10 µl aliquot of this bacterial suspension was deposited onto carbon-coated nickel grids, stained with 0.1% uranyl acetate, air-dried, and finally observed under a JEOL JEM1400 electron microscope.

### Quantitative real-time PCR

Wild-type cells of *F. nucleatum* containing the pZP4C(*ftsZ*) were initiated from a single colony and inoculated in a TSPC medium supplemented with 5 µg/ml thiamphenicol. Without the inducer theophylline, these cells were cultivated overnight at 37°C in an anaerobic chamber. Following the overnight growth, two subcultures were established at a 1:1000 dilution: one with 2 mM theophylline and the other without, serving as the control. After a 20-hour incubation, cells from both cultures were harvested by centrifugation. The harvested cell pellets were resuspended in Trizol (Ambion) and subjected to mechanical disruption using 0.1 mm silicon spheres (MP Bio) for lysis. Subsequent total RNA extraction was achieved with a Direct-zol RNA MiniPrep kit (Zymo Research), and the extracted RNA was reverse transcribed to cDNA using the SuperScript III reverse transcriptase (Invitrogen). For quantitative real-time PCR analysis, cDNA was mixed with iTAQ SYBR green supermix (Bio-Rad) and primer sets targeted to *ftsZ* and *ftsA* (Table 2). Gene expression levels were deduced using the 2^-ΔΔCt^ method (70), where the 16s rRNA gene acted as the normalization control. All procedures were consistently validated across 2 independent experiments, each conducted in triplicate.

### Indole assay and qualification

Two bacterial strains underwent an indole assay: the first group consisted of *F. periodonticum* wild-type (used as a control) and wild-type cells harboring the pZP4C(*tnaA_Fp_*) plasmid. Similarly, the second group included *F. nucleatum subsp. nucleatum* CTI-2 (serving as a control) and a wild-type cell harboring the pZP4C(*tnaA_CTI-2_*) plasmid. For strains with pZP4C plasmids, initial cultures were established overnight without an inducer and then diluted 1:1000 in fresh TSPC medium with various concentrations of theophylline (0, 0.1, and 2.0 mM), followed by a 22-hour anaerobic incubation at 37°C. After incubation, 200 µl samples from each culture were subjected to the indole test using Kovacs’s reagent. A positive test appeared as a pink to red hue, while the negative was yellow, with images captured 10 minutes post-reagent addition. The indole levels were quantified using a refined method we previously detailed. Briefly, supernatants from centrifuged cultures were tested with Kovacs reagent, and absorbance readings were matched to a standard curve for concentration determination.

## AUTHOR CONTRIBUTIONS

C.W. conceived and designed all experiments. P.Z., B.G., F.S., and C.W. performed all experiments. P.Z., B.G., and C. W. analyzed data. C.W. wrote the manuscript with contribution and approval from all authors.

## DATA AVAILABILITY STATEMENT

Materials are available upon reasonable request with a material transfer agreement with UTHealth for bacterial strains or plasmids. Plasmids information can be accessed on Benchling links at https://benchling.com/s/seq-BfSBxC6Ip4TX3HNnN0dX?m=slm-ImgpufGbZIscvdntMWhY (pZP4C) and https://benchling.com/s/seq-XynsExcpJvdQJxGszULy?m=slm-IH7R34mzlIQtZQ3ufOwk (pZP4C(*mCherry*)).

## ACKNOWLEDGMENTS

This research received funding from the National Institute of Dental and Craniofacial Research (NIDCR) under grant number DE030895, awarded to C.W. We extend our gratitude to Dr. David S. Weiss from the Department of Microbiology and Immunology at the University of Iowa for generously providing the pIA33 plasmid. Additionally, our thanks go to Dr. Wendy Garret of Harvard T.H. Chan School of Public Health for supplying the *F. nucleatum* CTI-2 strain sourced from CRC samples.

